# Interactions of Elongated Dinuclear Metallo-Cylinders with DNA Three-Way and Four-Way Junctions

**DOI:** 10.1101/2025.07.06.663355

**Authors:** Samuel J. Dettmer, Hannah M. P. Stock, Michael J. Hannon

## Abstract

Non-canonical DNA structures play important roles in processing of the genetic code. Three-way (3WJ) and four-way (4WJ) junctions are dynamic, multi-stranded structures containing an open cavity at the centre. We have previously demonstrated that supramolecular dinuclear metallo-cylinders bind well inside 3WJ cavities, having an optimally complementary size and shape match, cationic charge to bind the anion, as well as the ability to π-stack with the branchpoint nucleobases. Herein we show that a longer metallo-cylinder with a similar but extended central π-surface, binds to both 3WJ and 4WJ structures with good selectivity over double-stranded DNA. Experimental investigations, informed by molecular dynamics (MD) simulations, reveal that while this longer cylinder can bind 3WJs as the previously studied cylinders, the extended π-surface of the cylinder now also facilitates 4WJ binding. The simulations capture two metastable 4WJ conformations – one resembling a 3WJ, and another where the extended length enables the cylinder to angle into and stabilise a rhombus-shaped 4WJ cavity. The ability to tune the structure of supramolecular assemblies is important for targeting different DNA structures with varying specificity and in this work we demonstrate the usefulness of overall length as a parameter for modulating DNA binding.

## Introduction

Deoxyribonucleic acid (DNA) is an abundant and important biomolecule that stores genetic information. Though primarily found in the double helix structure, in order for the genetic information to be processed, the helix must unwind and adopt dynamic conformations, which allow proteins to bind and perform tasks [1-3]. Such non-canonical structures include forks, three-way junctions (3WJ) and four-way junctions (4WJ), which offer binding cavities for nano-sized compounds and are potential therapeutic targets for diseases associated with these structures [4-8]. Current DNA-targeting cancer drugs in the clinic act by coordinating to the purine bases, or intercalating between the base pairs in duplex DNA, which prevents DNA replication and cell division, and leads to cell death [9-11]. Whilst effective, these drugs pose two primary issues. Firstly, they do not discriminate between healthy and cancerous cells. Secondly, these drugs bind non-selectively to duplex DNA and thus affect many different genes and cellular processes. These limitations, together with off-target protein interactions, lead to significant side-effects and result in the patient experiencing significant discomfort. Targeting non-canonical structures such as DNA junctions offers an attractive route both to gaining greater specificity and reducing side-effects.

We have shown previously that metallo-supramolecular assembly allows us to construct large nano-size structures that are new types of drugs with excellent 3D shape match for DNA junctions. In particular X-ray crystallographic and solution NMR structures reveal that a class of dinuclear triple helicate metallo-cylinders (∼2 nm in length and ∼1 nm in diameter) of formula [M_2_(L1)_3_]^4+^ are a perfect match for the trigonal prismatic cavity of a DNA 3WJ (*Fig. 1A*) [12-16]. The central core of these cylinders bears outward-facing aromatic surfaces, allowing them to thread beautifully into the cavity of the 3WJ and for the aromatic faces to interact with the central by π-π stacking. This has stimulated other teams to also design supramolecular compounds to bind 3WJs and explore biological activity [17-37].

**Figure 1.**
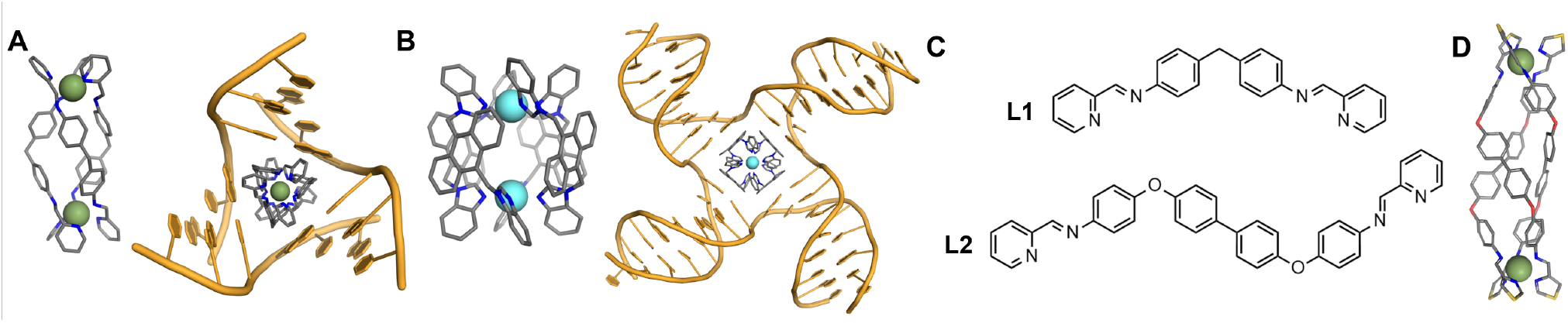
A) 3D model of the Fe_2_(L1)_3_ cylinder and the crystal structure of this cylinder bound inside a 3WJ [12]. B) 3D model of a dinuclear platinum cage and an MD snapshot of this compound bound inside a 4WJ [43]. C) Structures of ligands L1 and L2. D) Crystal structure of the iron triple-helicate reported by Howard-Smith *et al*. using thiazoleimine ligands [46].

4WJs are analogous structures comprised of 4 helical domains that meet at a branchpoint. These structures are prevalent in biology, occurring across the genome as a major intermediate in homologous recombination – a process integral to DNA repair [38] as well as genetic diversity. They are also found in areas of the genome that contain inverted repeat sequences, where they are associated with a variety of diseases. 4WJs are known to adopt two primary conformations – open (cruciform) and closed stacked (stacked X-shape) [39-41]. The open conformation is the biologically relevant conformation, often stabilised by proteins, and we have previously shown that metallo-compounds displaying square symmetry stabilise the 4WJ by binding inside the open cavity (*Fig. 1B*) [42, 43]

Dinuclear metallo-supramolecular compounds are also of interest in the field of materials chemistry due to their spin crossover properties [44, 45], and in this context Howard-Smith *et al*. recently reported long triple-helicates (*Fig. 1D*) (∼3 nm in length) [46]. Other than their increased length, the width and general twisted-prismatic shape of this helicate is broadly the same as the metallo-cylinder we developed [12]. The key difference lies in the central core of the ligand which is elongated from a diphenylmethane in L1 to a 4,4’-bis(phenoxy)biphenyl. This endows the central section of the helicate with more extensive *π*-surfaces presented as the shell of the cylinder structure. We were interested to interrogate what effect elongating the *π*-surfaces would have on the DNA-junction binding behaviour, as part of our wider quest to access novel types of DNA structure recognition.

## Results and Discussion

### Synthesis and Characterisation

Howard-Smith *et al*. described elongated helicates based on thiazoleimine ligands [46]; for our designs we prepared analogous ligands using pyridylimine binding units. The ligand L2 was synthesised by the condensation of 2-pyridinecarboxaldehyde and 4,4′-(1,1′-biphenyl-4,4′-diyldioxy)dianiline in ethanol in the presence of acetic acid as catalyst and isolated by filtration in 80% yield (*Figs. S1-S2*). The iron(II) and nickel(II) helicates were then synthesised by mixing ligand L2 in a 3:2 ratio with either iron(II) tetrafluoroborate or nickel(II) tetrafluoroborate in acetonitrile and warming. In each case, the compounds were characterised by NMR (though the Ni compound is paramagnetic) and electrospray ionisation mass spectrometry (ESI-MS) revealed a dominant peak corresponding to [M_2_(L2)_3_]^4+^, consistent with a dinuclear triple-stranded species in solution (*Figs. S3-S8*).

The aqueous solubility of the tetrafluoroborate salts of these complexes was poor and not suitable for biophysical experiments, so the compounds were obtained as acetate salts by reacting either iron(II) acetate or nickel(II) acetate with the ligand in methanol to form [Fe_2_(L2)_3_](OAc)_4_ and [Ni_2_(L2)_3_](OAc)_4_. These salts demonstrated much better solubility in water and their [M_2_(L2)_3_]^4+^ stoichiometries were again confirmed by ESI mass spectrometry (*Fig S9*.). The Fe cylinder proved to be unstable in solutions containing sodium salts (NaCl, NaOAc, NaNO_3_; *Fig. S10*; the presence of sodium cations is important for biophysical DNA studies), evidenced by a change in solution colour from purple to colourless over the course of 1 hour. The Ni compound also showed some instability in buffer conditions (89 mM Tris, 89 mM boric acid, 50 Mm NaCl) over 12 hours, evidenced by changes in the UV-VIS spectrum (*Fig. S11*), though ESI-MS confirmed that the cylinder was still the major species (*Fig. S12*). The second largest peak is attributed to the 4,4′-(1,1′-biphenyl-4,4′-diyldioxy)dianiline spacer unit – the product of hydrolysis at the ligand imine bonds – and M_2_L_2_ type species were also detected in the baseline. With this in mind, we proceeded with using this nickel(II) cylinder for our biophysical studies, using freshly dissolved compound each time.

Interestingly, the ^1^H NMR of the diamagnetic iron(II) (low spin) complex revealed the presence of two species present in solution in a ratio of approximately 3:1 (*Fig. 2A*). In our original triple-helicate metallo-cylinders the diphenylmethane spacer constrains flexibility in the helicate and triple-helical *rac-*isomers (*ΛΛ* and *ΔΔ*) result. However we have previously shown that when we extend that spacer slightly (adding two additional imines) a *meso-*isomer (mesocate: *ΛΔ*) can also be formed and it appears that this is also the case for this triple-stranded complex [47]. During the course of our study, a solid-state crystal structure of this complex has appeared, revealing a *rac-*helicate structure in the solid-state. However, in that publication the presence of the different isomers is not recognised [48].

**Figure 2.**
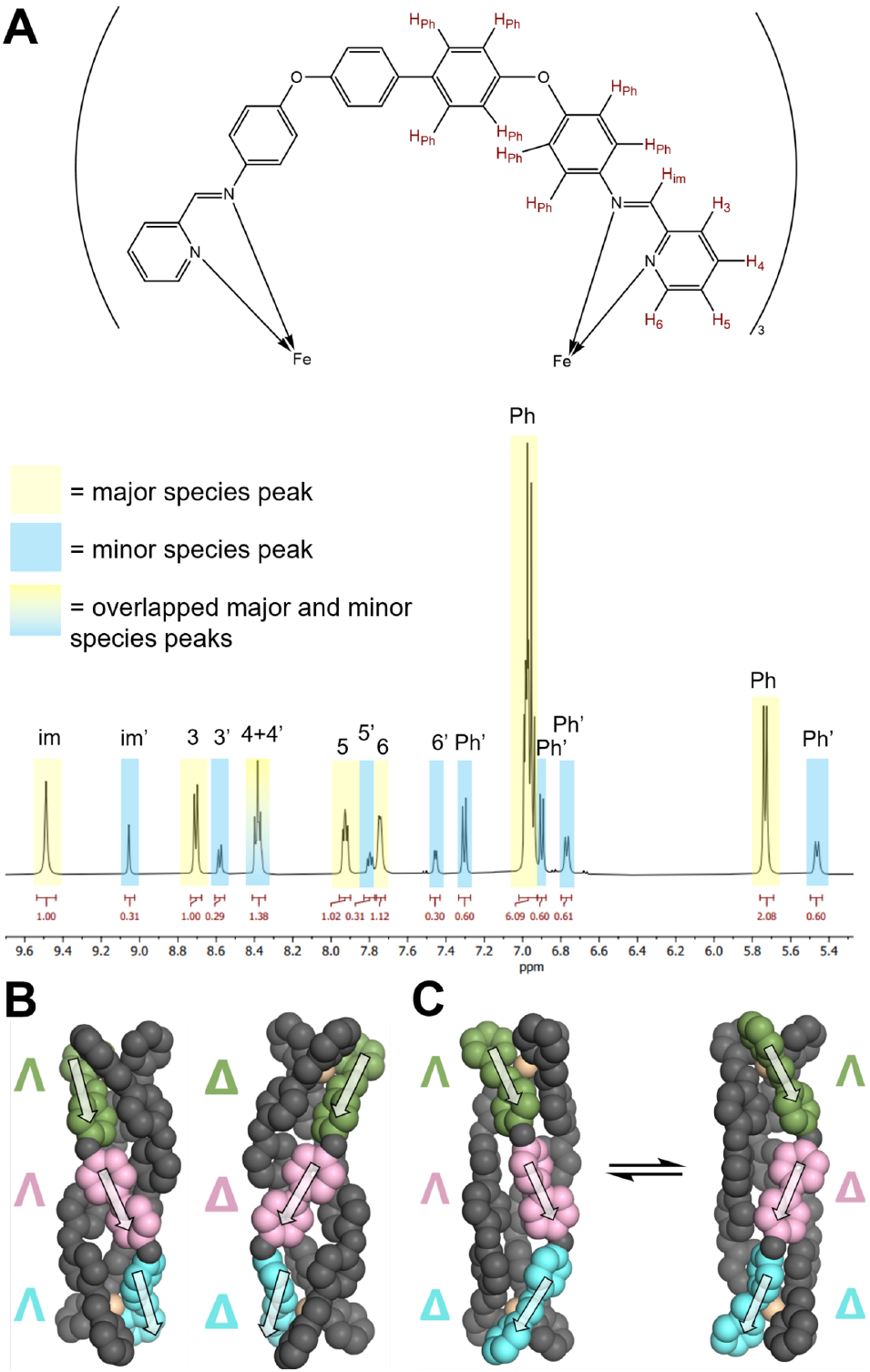
A) NMR spectrum of the [Fe_2_(L2)_3_](BF_4_)_4_ cylinder recorded in acetonitrile, zoomed in on the region 5.4-9.6 ppm, where the complex peaks appear. Highlighted in yellow are peaks corresponding to the major species, highlighted in blue are peaks corresponding to the minor species and highlighted in the yellow/blue gradient (peak at approx. 8.4 ppm) are overlapped major/minor species peaks. B) 3D DFT models of the two enantiomers of the helicate structure and C) the ∧∧Δ mesocate interconversion with its ∧ΔΔ enantiomer, as seen in MD. The green, pink and cyan atoms with overlaid arrows are to aid visualisation of the three positions containing chiral information.

In order to explore the idea of a mesocate isomer further, we employed DFT geometry optimisation calculations. Firstly, optimisation of the helicate was performed using the reported crystal structure as an initial guess. This generated a model containing the key structural features seen in the solid state, albeit with a slightly longer metal-metal distance (19.7 Å vs 18.7 Å). From this structure, a mesocate isomer was generated in the Avogadro molecular editor [49], by separating the two halves of the helicate at the biphenyl linker and inverting the chirality of one half before rejoining the bonds and energy minimising. The resulting structure was then fed into a DFT geometry optimisation.

In our original, smaller (L1) cylinder, local chirality is found at each of the two metal centres (either Λ or Δ). The observed helicate structure has the same chirality at each metal leading to two enantiomers: ΛΛ and ΔΔ. A hypothetical mesocate would have different chirality at each metal (and be designated ΛΔ); this isomer is not observed for the L1 cylinder. With the elongated spacer unit of the L2 cylinder, local chirality can also be found in the biphenyl unit, which could orient its axis in either Λ or Δ directions, independent of the metal centres, due to flexibility at the oxygen atoms. In the helicate crystal structure, the ligands twist in the same direction along the length of the compound, giving rise to a ΛΛΛ helicate (and a corresponding ΔΔΔ enantiomer; *Fig. 2B*). The DFT optimised mesocate structure was generated from the DFT model of the ΔΔΔ helicate and thus has a Δ twist at the biphenyl, giving rise to a ΛΔΔ cylinder. This is not a true meso-isomer, as a ΛΛΔ cylinder would be its enantiomer.

The DFT calculations were used to generate molecular dynamics (MD) parameters for the compounds and the two isomers were each simulated alone in presence of water (with Cl^-^ counterions). Over the course of 3 µs, two simulations captured the interconversion of the ΔΔΛ mesocate to its ΔΛΛ enantiomer (*Fig. 2C*) by concerted rotation of the two metal centres. Rapid interconversion between the two enantiomers is consistent with the observation of only two species in NMR, whereby the ΛΛΔ and ΔΔΛ mesocates interconvert faster than the NMR timescale. This suggests it may also be possible that the ΛΛΛ and ΔΔΔ helicates also rapidly interconvert with their respective ΛΔΛ and ΔΛΔ isomers, though this was not observed within the timescale of MD (5 µs).

### Three-Way Junction Binding

To investigate whether this larger helicate binds to DNA 3WJs, we employed polyacrylamide gel electrophoresis (PAGE; *Fig. 3*). Following the established 3WJ-binding assay protocol, we utilised three individual DNA oligomers bearing appropriate sequences to form a 3WJ and exposed them to various concentrations of the water soluble [Ni_2_(L2)_3_](OAc)_4_ cylinder [15]. In the absence of a suitable binding agent, at room temperature these strands do not assemble into a larger structure and remain single stranded, whereas in the presence of suitable binding agents, the 3WJ is stabilised and a distinct new slower-migrating band appears on the gel. This offers an on/off visualisation of 3WJ binding and can be seen in Fig. 3 (lane 1 vs lane 2) for the positive control [Ni_2_(L1)_3_]^4+^. The elongated cylinder [Ni_2_(L2)_3_](OAc)_4_ also binds and stabilises the 3WJ (lanes 4-7) and does so in a concentration dependent fashion. However, the intensity of the 3WJ band is noticeably lower than in the positive control, particularly at 0.5 and 1 equivalents. This may signify that the binding is weaker than for the L1 cylinder due to the lower charge density, although given that the compound does degrade to some extent in solution (*Figs. S11-12*), it is possible that the lower intensity may reflect this (likely it is a combination of both effects).

**Figure 3.**
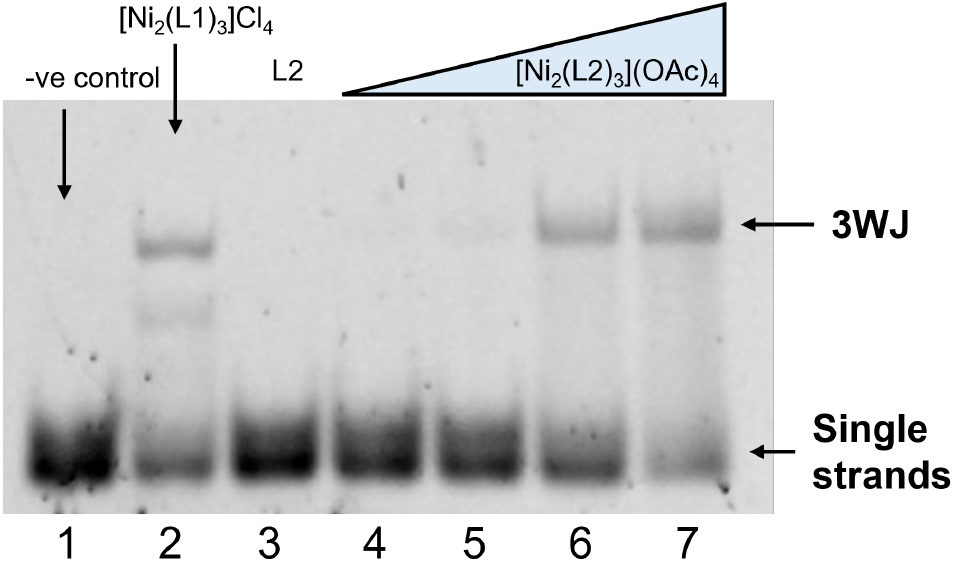
PAGE gel showing binding of the cylinder to the 3WJ. Lane 1 contains 3WJ alone, lane 2 contains the 3WJ + Ni L1cylinder, lane 3 contains the free ligand, lanes 4-7 contain 3WJ + 0.5, 1, 2 and 4 equivalents of the Ni cylinder. Samples contained 1X TB running buffer and 50 mM NaCl

The observed 3WJ band with [Ni_2_(L2)_3_]^4+^ is slightly retarded with respect to that observed with the [Ni_2_(L1)_3_]^4+^ cylinder positive control, and this gel shift indicates a greater hydrodynamic radius for the 3WJ-cylinder complex which is expected given the larger dimensions of this elongated cylinder, and consistent with a section of the cylinder protruding out from the 3WJ cavity. The absence of a 3WJ band when the DNA strands are exposed to just the ligand L2, confirms that it is the metal complex that induces 3WJ formation and not solely an organic ligand component. Interestingly, at this concentration of NaCl (50 mM), the L1 cylinder also weakly induces formation of a two-stranded fork structure, whereas the L2 cylinder does not, suggesting that the L2 cylinder may have a lower affinity for Y-fork two-stranded structures than the L1 cylinder.

Molecular dynamics (MD) simulations have proved valuable tools for providing insight into DNA junction binding interactions at the atomic level and we have successfully used them previously to help explain experimental observations [42] and screen compounds for their junction binding fit [43]. We therefore performed molecular dynamics (MD) simulations here to probe more deeply the binding of the elongated cylinder.

Using a 3WJ structure adapted from PDB 1F44 [50], MD simulations of the DFT optimised Ni L2 helicate with the 3WJ were initially employed. Six simulations, where the helicate was placed outside (but close to) the cavity, were first run (3 per enantiomer). Only two of these captured the entry of the helicate into the 3WJ cavity but once the helicate entered it bound stably over the timescale of the simulation (1 µs; *Fig 4A*). In the other four simulations, the 3WJ cavity closed up (allowing two of the arms to stack coaxially) before the helicate could reach the cavity and thus did not capture this binding mode. In these cases, the helicate instead bound along, or at the end of, the duplex arms (*Fig. S13*) which is valuable as a model of how they will engage with duplex DNA. In terms of the 3WJ closing before cylinder entry, it is worth noting that this larger L2 cylinder will move more slowly in MD than the short L1 cylinder because of its higher mass – and so enter more slowly from the same starting position - and that the timescale of re-opening of junction structures is too slow to be captured in MD simulations [51]. This is not an issue experimentally in the gel studies as junction structures are dynamic and timescales longer, and indeed the short DNAs used will be exchanging between 3WJ (minor) and single stranded (major) configurations prior to cylinder addition.

**Figure 4.**
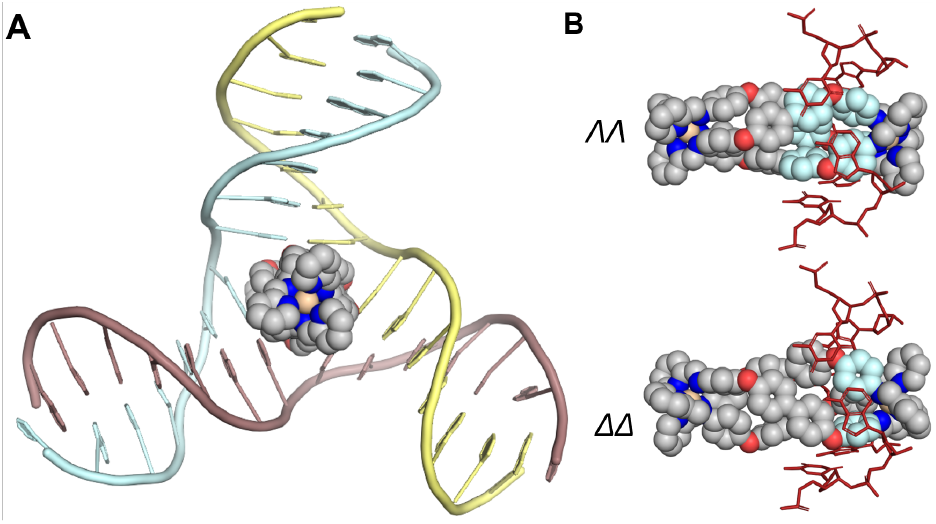
A) MD snapshot of the [Ni_2_(L2)_3_] _4+_helicate (M enantiomer - *∧∧*) bound in the cavity of a 3WJ. B) Close-up image of the two enantiomers of the cylinder bound inside the 3WJ. The aromatic surfaces involved in face-face π stacking, and in CH-TX stacking are coloured in light blue. Only the branchpoint DNA bases are shown. Hydrogens have been omitted in all images for clarity.

Once the ability to enter the 3WJ in MD was confirmed, a further six simulations (3 per enantiomer, 1 µs each) were undertaken with the [Ni_2_(L2)_3_]^4+^ helicate starting in the bound cavity position; in each case the enantiomers remained bound inside the junction for the duration of the simulations (*Fig. 4A*). In the case of the *ΛΛ* enantiomer, each branchpoint base pair is engaged in π-stacking interactions with one of the biphenyl rings and a phenoxy ring on an adjacent ligand, causing these two rings to orient themselves coplanar with each other to facilitate this interaction (*Fig. 4B*). This interaction is very similar to the crystallographically characterised structure for the [Ni_2_(L1)_3_]^4+^ *ΛΛ* enantiomer complex with a DNA 3WJ (*Figs. 1A, S14)*. The rest of the [Ni_2_(L2)_3_]^4+^ complex protrudes out of the 3WJ, and the simulations do not reveal a strong preference for which side of the junction (where the major or minor grooves meet) the cylinder protrudes from (both are observed). The phenyl rings of the spacer that are π-stacked with the DNA bases in the 3WJ cavity have only restricted motion, while those that protrude into solution spin more freely. The *P* (*ΔΔ*) enantiomer has less overlap between the aryl faces and the branchpoint base pairs but is nevertheless still held tightly in place with the aid of π interactions (*Fig. 4B*) and is otherwise very similar in its binding. Simulations with the Fe variants of the cylinder yielded the same observations.

Equivalent simulations were run, in which the mesocate isomers were placed inside the junction (3 per enantiomer at 1 µs each). As seen with the helicate, only one end of the mesocate interacted with the nucleobases at any one time, when bound to a fully base-paired 3WJ. In all simulations with the ΛΔΔ isomer, this was the Δ end and the binding interactions were very similar to those seen with the ΔΔΔ helicate (*Fig. S16A*). Simulations with the ΛΛΔ isomer captured a more varied conformational landscape. One simulation showed this isomer binding with its Λ end, analogous to the ΛΛΛ helicate. One simulation captured a metastable conformation in which a base pair was broken and those bases interacted separately with the cylinder (*Fig. S15*). In the third simulation, interconversion of the ΛΛΔ mesocate to its ΛΔΔ enantiomer was observed, and the cylinder proceeded to bind with the Δ end. These simulations suggest that the binding end of the mesocate is governed by the twist of the biphenyl, and the end bearing the matching metal centre chirality is preferred for binding.

A further 3 simulations were run where the ΛΔΔ mesocate was placed with the Λ end inside the junction cavity. In 2 out of 3 simulations, the mesocate adopted a stable binding mode with similar interactions to the ΛΛ helicate, though the geometry of this isomer meant that the coplanarity of the binding phenyl rings was suboptimal (*Fig. S16A*). In the other simulation, the helicate initially bound with its Λ end but then shifted through the cavity such that the Δ end was binding (*Fig. S16B*).

### Four-Way Junction Binding

A similar PAGE protocol as for the 3WJ was first utilised to investigate the binding of the [Ni_2_(L2)_3_]^4+^ cylinder to 4WJs. This time, four individual strands with a 4WJ-forming design, which similarly does not assemble spontaneously unless a binding agent is present, were used. A distinct, new slower-migrating band appeared on treatment with [Ni_2_(L2)_3_]^4+^ (again faintly at low loading) corresponding to a 4WJ species (*Fig. 5A*). Additionally, smearing is visible between the 4WJ and single strand bands, indicating a (much) weaker binding to intermediate (two- or three-stranded) structures. The smaller [Ni_2_(L1)_3_]^4+^ cylinder, also forms the 4WJ band with a higher intensity, indicating strong binding to the 4WJ structure. However [Ni_2_(L1)_3_]^4+^ also stabilises a three-stranded “pseudo-3WJ” structure in the gel – a band not seen with the L2 cylinder. This may indicate that this structure is disfavoured for an elongated cylinder, though the overall binding is weaker across the board.

**Figure 5.**
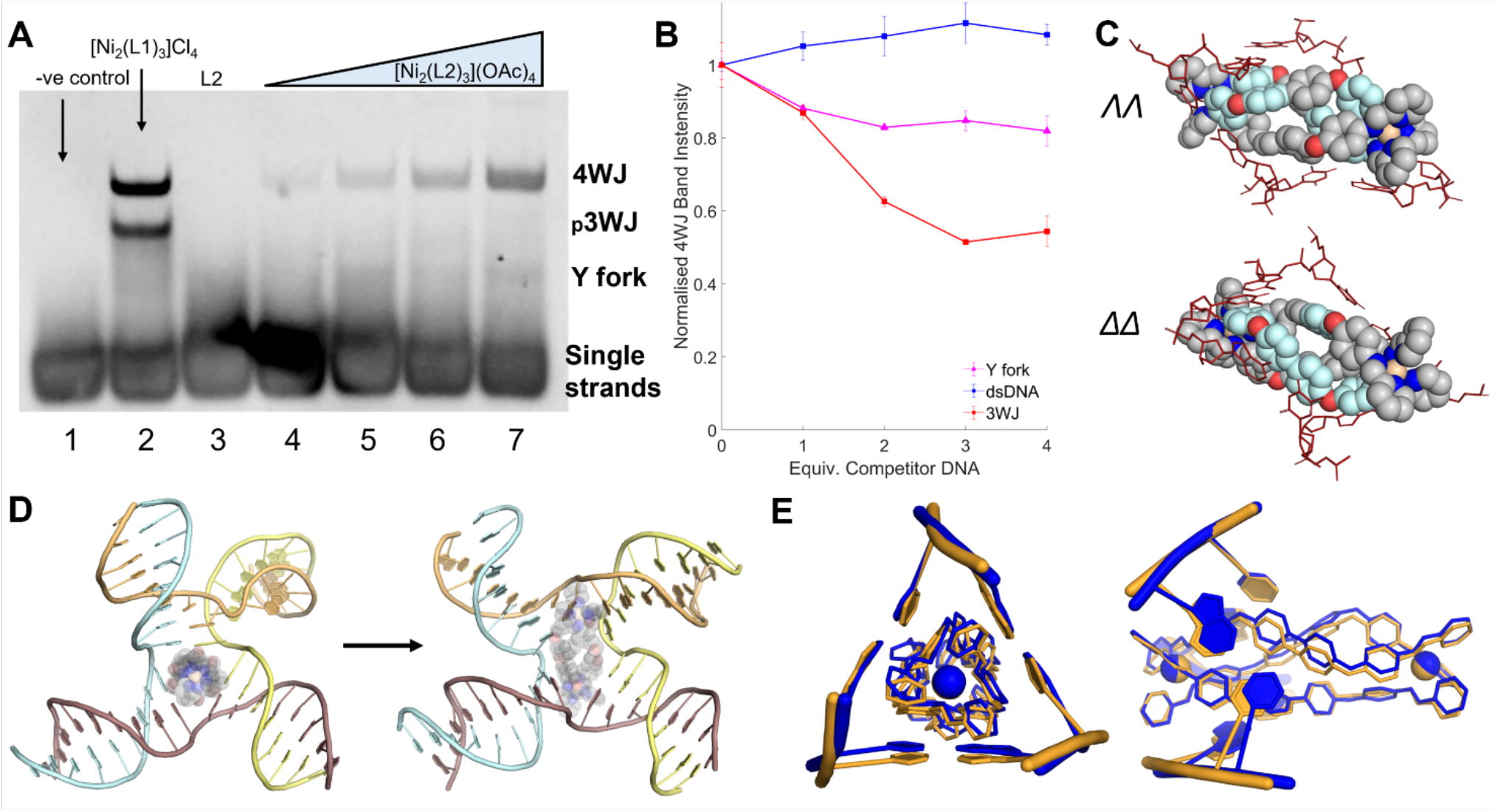
A) PAGE gel showing binding of the cylinder to the 4WJ. Lane 1 contains 4WJ alone, lane 2 contains the 4WJ + Ni L1 cylinder, lane 3 contains the free ligand, lanes 4-7 contain 4WJ + 0.5, 1, 2 and 4 equivalents of the Ni cylinder. Samples contained 1X TB running buffer and 10 mM NaCl. B) Normalised intensity curves of the PAGE competition gels. Competitor Y fork (magenta), dsDNA (blue) and 3WJ (red) were titrated into a fixed concentration of FAM-4WJ + 3 equiv. Ni L2 cylinder. Each point represents the normalised % intensity of the 4WJ band. Each point is the average of 3 repeats and error bars represent the standard deviation. C) Close-up image of the two enantiomers of the Ni L2 helicate isomer bound inside the 3WJ. The aromatic surfaces involved in face-face pi stacking are coloured in light blue. Only the branchpoint DNA bases are shown (in red). D) MD snapshots of the ∧∧ Ni L2 cylinder bound in a 3WJ-like 4WJ (left) and in a rhombus-shaped cavity 4WJ (right). The 3WJ-like structure is facilitated by the formation of a base pair between 2 non-adjacent strands (here the yellow and blue strands). The helicate has been made translucent to aid visualisation of the bases. The arrow denotes that the transition between these conformations was observed within a simulation (1 μs). E) Front-on and side-on MD snapshot overlays of the mesocate isomer bound inside the 3WJ (orange) and the 3WJ-like 4WJ (blue). Only the bases directly interacting with the mesocate are shown. Hydrogens have been omitted in all images for clarity.

**Figure 5.**
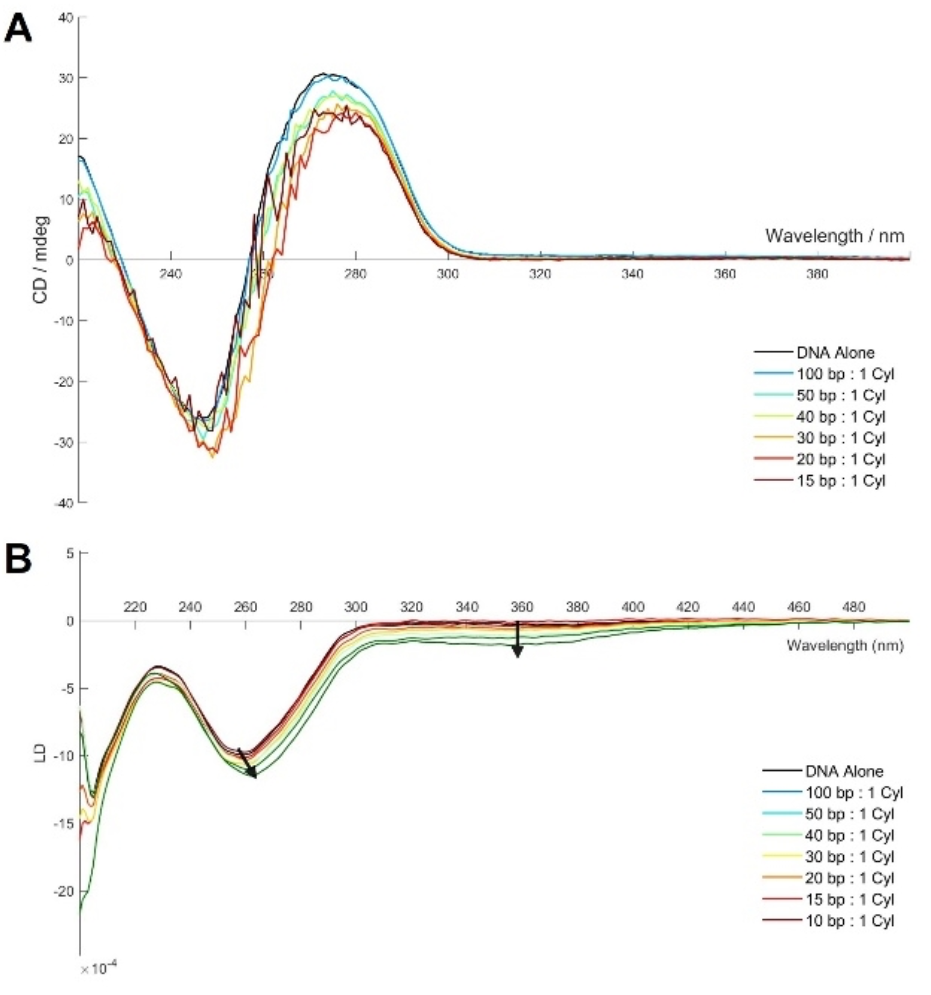
CD (A) and LD (B) spectrum of ctDNA (200 μM base pairs) with increasing loadings of [Ni_2_(L2)_3_](OAc)_4_ in 10 mM Na cacodylate, 100mM NaCl.

PAGE competition assays were employed to assess the preference of the compound for different DNA structures (*Fig. 5B*). The 4WJ was mixed with buffer and [Ni_2_(L2)_3_]^4+^ cylinder at constant concentration: 3 equivalents of cylinder were used to ensure a sufficiently strong 4WJ signal. This was then mixed with increasing amounts of competitor DNA (3WJ and dsDNA) and the intensity of the 4WJ PAGE bands subsequently measured. A fluorescently labelled (FAM) 4WJ was utilised to prevent noise from potential uneven staining of the gel and to leave the competing DNA structures silent. As the competitor DNA structures bind with the cylinder, this alters the amount of free cylinder in solution and so the dynamic equilibrium leads to a decrease in the 4WJ band (*Fig. S17*).

The results demonstrate that dsDNA is not a competitor at these ratios, with little to no change (within error) in the 4WJ band. On the other hand, the 3WJ does indeed bind to the compound and cause a decrease in the 4WJ band of ∼50% at 3 equivalents (relative to the 4WJ, 1 equivalent relative to cylinder). The Y fork structure also competes, albeit less so than the 3WJ, reducing the intensity to ∼85% at 3 equivalents. Similar competition experiments were attempted using a FAM-labelled 3WJ and titrating in 4WJ, however the initial 3WJ intensity was too low to obtain reliable results.

MD simulations were again employed to investigate the interaction with the 4WJ. Simulations were initially started with the [Ni_2_(L2)_3_]^4+^ helicate inside the open 4WJ cavity (3 per enantiomer, 1 µs each). The majority (5 out of 6) of simulations settled into a metastable state in which the helicate is angled relative to the axis of the 4WJ cavity – this binding is unlike that observed with the 3WJ, where the helicate threaded perfectly through the centre of the cavity. This binding mode leads to a rhombic-shaped cavity (*Figs. 5C, 5D*), more reminiscent of the peptide-bound 4WJ structure reported by Ghosh and colleagues [40], as opposed to the square shaped cavity seen in simulations with the pillarplexes and anthracene cage compounds [42, 43]. This binding mode is enabled by the length of the cylinder, and it allows the aromatic faces across the full span of the ligands to interact with all four branchpoint base pairs (*Fig. 5C*). One simulation captured a much more dynamic, half-closed 4WJ state, wherein the cylinder was bound at one side of the cavity, leaving the other side free to close up and coaxially stack those duplex arms (*Fig. S18*). Simulations, in which the helicate’s initial position was outside the junction were not able to consistently capture the entry of the helicate into the junction, before the 4WJ transitioned to the closed form (for the same reasons as in the 3WJ simulation and the same issue of timescales of re-opening). Again, simulations with the Fe variants of the cylinder yielded similar observations of the rhombus shaped 4WJ cavity.

Simulations with both mesocate isomers were also performed (3 per enantiomer, 1 µs each) and most often captured a much more 3WJ-like structure, in which a triangular shaped cavity formed around the helicate. The mesocate was able to stabilise an AT Watson-Crick pairing across 2 non-adjacent strands (*Fig. 5D*), completing the branchpoint base pairing of a 3WJ and effectively excluding one of the 4WJ strands from interaction with the mesocate (*Fig. 5E*). The rhombic-shaped cavity conformation was observed in one simulation, and transient conversion between the conformations was also observed in another, indicating that both conformations are accessible.

As MD simulations operate on a very small timescale in comparison to experimental conditions, they can be influenced by the initial starting position of the DNA and binder. To mitigate against bias here, further simulations were run, starting the compound in the less-observed conformation for that isomer (helicate in the 3WJ-like 4WJ, and ΛΔΔ mesocate in the rhombic-cavity 4WJ). In all 3 simulations from this starting position, the ΔΔ helicate remained stably bound to the 3WJ-like structure. As was observed with the mesocate, the new branchpoint base pair arising between non-adjacent strands was maintained, and the helicate engaged in near identical interactions to those seen with the 3WJ itself. This was also observed in one of the simulations of the ΛΛ helicate in the 3WJ-like structure, however the other 2 simulations captured the conversion of the 3WJ-like structure into the rhombic-cavity structure (*Fig. 5D*), with the helicate adopting a similar binding position as in the original 4WJ simulations, where this conformation was first observed. This transition is enabled by the breaking of the AT base pair between the non-adjacent strands,which then allows the 4WJ to reassemble its “true” branchpoint base pairing.

In 2 out of 3 simulations where the mesocate was place inside the rhombus cavity, the mesocate adopted a similar binding mode as the helicate did, though it is not able to make full use of its *π* surface capacity due to the twist of the ligands in this isomer (*Fig. S19*). In the other simulation, the system was much more dynamic and transformed away from the rhombic cavity conformation, towards a 3WJ-like conformation, though the metastable conformation with the 3 base pairs intact around the cylinder was not achieved. These additional simulations thus give greater confidence that the mesocate and helicates do adequately explore the energy landscape in the timescale of the simulations and that the differences in their binding are reproduced from different starting positions.

### Double Stranded DNA Binding

Whilst junctions form dynamically during processing, DNA is stored, and spends most of it time, as double-stranded DNA (dsDNA) in the well-known double helix structure. Although the original [M_2_(L1)_3_]^4+^ cylinders bind preferentially to 3WJ they also bind (more weakly) to double stranded DNA and cause intramolecular coiling [52]. The competition experiments herein show that the elongated [M_2_(L2)_3_]^4+^ cylinders also strongly prefer junctions over dsDNA but we were interested to investigate whether the increased length of this helicate would lead to any differences in DNA coiling behaviour.

Circular dichroism (CD) and linear dichroism (LD) are valuable tools for investigating changes in DNA conformation. CD explores the chiral nature of the DNA helix, whilst LD uses a shear force to orient the DNA along a rotational flow. Calf-thymus DNA (ctDNA) was utilised as an in vitro analogue for genomic DNA and was made up as a 1 mM (in base pairs; concentration confirmed by UV-VIS spectroscopy using ε = 13100 M cm^-1^ bp^-1^) stock solution in water. This was then diluted to 200 µM in buffer (10 mM Na cacodylate and 100 mM NaCl) and to this, increasing amounts of [Ni_2_(L2)_3_](OAc)_4_ cylinder were added and the CD and LD spectra were recorded.

ctDNA exhibits a characteristic CD curve with a positive peak at 280 nm and a negative peak at 245 nm. Upon increasing additions of the [Ni_2_(L2)_3_]^4+^ cylinder, there is little change in the negative peak, but the intensity of the positive peak decreases with a slight red shift (*Fig. 6A*). Notably, the curves become increasingly more noisy at higher loadings. The overall profile of the spectrum retains the characteristics of ctDNA, indicating that the cylinder is not having a significant effect on the basic B-form DNA structure. The observed noise at higher loadings might reflect some aggregation promoted by charge neutralisation between the negatively charged DNA phosphate backbone and the tetracationic cylinders, leading to scattering by the resultant particles.

In the LD experiment the DNA is oriented in flow in a Couette cell, and the orientation probed by difference in absorption of light polarised parallel and perpendicular to the direction of flow. ctDNA exhibits a characteristic negative peak at 260 nm in its LD spectrum. When the smaller [Ni_2_(L1)_3_]^4+^ cylinder is added it coils the ct-DNA leading to a rapid loss of that 260 nm band [52]. By contrast, titration of the [Ni_2_(L2)_3_]^4+^ cylinder into ctDNA (200 µM in base pairs, 10 mM Na cacodylate, 100 mM NaCl) saw a slight increase and slight red shift in the intensity of the 260 nm peak, and the emergence of a broad band between 300 and 400 nm (*Fig. 6B*). This new induced band roughly aligns with the absorbance spectrum of the cylinder, indicating both that the cylinder is interacting with the ctDNA, and that it is doing so in an oriented manner – most likely by sitting in the major or minor groove. However, unlike the parent L1 cylinder, binding of the L2 cylinder does not cause coiling or unwinding of the ctDNA.

MD simulations of both isomers with a short 25mer dsDNA capture metastable biding at the termini of the duplex and in the minor groove (*Fig. S20*). In some cases, the base pairing at the ends of the duplex were frayed open and the cylinder resided within this opened end, though the ctDNA used in the experimental studies, and by extension genomic DNA, contain far fewer termini so this binding mode will be less prevalent with these DNA systems. The minor groove binding is consistent with the presence of an induced LD band.

## Conclusion

Herein we have described a longer metallo-cylinder and shown that it binds to both 3WJs and 4WJs, and does so in preference to binding to dsDNA. Simulations show that increased length of the L2 cylinder compared with the previously reported L1 cylinder does not modify the type of binding to 3WJs, as it binds using only one end. The binding moiety on the L2 cylinder (in both helicate and mesocate forms) is topographically identical to the diphenylmethylene spacer unit of the L1 cylinder, leading to identical π-stacking interactions.

By contrast, the length of the L2 cylinder appears to be beneficial for binding 4WJs. Whereas the L1 cylinder tends to displace a strand from the 4WJ to form a pseudo-3WJ where it can bind, this is not seen for the L2 cylinder. Simulations show that the greater available π-surfaces of this elongated cylinder allow it to angle itself relative to the cavity, allowing for π-stacking interactions to occur along the entire length of the cylinder. Because the whole cylinder engages with the 4WJ, the simulations imply slightly different binding modes for the helicate and mesocate forms. The experimental and simulation results suggest that by increasing the π-surfaces, larger junction structures can be targeted. Agents that bind and potentially interfere with multiple different junction structures can have powerful biological effects [25].

Our experimental competition assays reveal a strong preference for junctions over double stranded DNA. Intriguingly, the L2 cylinder does not coil, or cause conformational changes in dsDNA in the same way that the L1 cylinder does, suggesting that the smaller size of that cylinder is important in its ability to condense DNA. While this [Ni_2_(L2)_3_]^4+^ cylinder has provided valuable insight into the design of future junction binders, the precise spacer used in this design does not provide sufficient mechanical coupling between the two metal centres and so two isomers arise which complicates their study. Combined with the issues of solution stability this means that they do not merit further study in cells. But nevertheless these in vitro biophysical studies provide a starting point for optimising these compounds and show that the length of the compound and of their available π-surfaces is another parameter that can be used for fine tuning the DNA binding of supramolecular assemblies.

## Acknowledgments and Funding

This work was funded by the BBSRC Midlands Integrative Biosciences Training Partnership (BB/T00746X/1), and the University of Birmingham. The authors acknowledge use of the University of Birmingham Analytical Chemistry facilities. All simulations were performed using the University of Birmingham BlueBEAR and CaStLeS HPC facilities. The authors thank Dr Catherine Hooper (U. Birmingham) and Dr Larry Melidis (U. Cambridge) for helpful discussions and advice.

## Supplementary Information and Data

### Availability

Supplementary material comprises supporting data and experimental methods. The authors have included additional references in the supplementary information. [53-61]. The data for the experiments contained in this manuscript together with videos of the simulations described are available via the UBIRA repository at https://doi.org/10.25500/edata.bham.00001275.

## Author contributions

MJH conceived the project and supervised and directed the studies. SJD led, and with HMPS, undertook gel experiments and MD simulations. SJD undertook CD and LD spectroscopy experiments and DFT calculations. HMPS undertook synthesis, compound characterisation and UV-VIS spectroscopy. All three authors drafted the manuscript.

