## Supplementary Information for "Interactions of Elongated Dinuclear Metallo-Cylinders with DNA Three-Way and Four-Way Junctions"

Electronic Supplementary Information

In addition to the Supplementary Information herein, all original NMR, mass spectrometry, gel electrophoresis, UV-Vis, CD, LD and simulation data, as well as videos for the simulations, are also made available online at: https://doi.org/10.25500/edata.bham.00001275.

### TABLE OF CONTENTS

Supplementary Data 2
- **Figure S1.** NMR spectrum of L2 ligand 2
- **Figure S2.** Mass spectrum of L2 ligand 2
- **Figure S3.** NMR spectrum of $[Fe_2(L2)_3](BF_4)_4$ 3
- **Figure S4.** COSY spectrum of $[Fe_2(L2)_3](BF_4)_4$ 3
- **Figure S5.** NOESY spectrum of $[Fe_2(L2)_3](BF_4)_4$ 4
- **Figure S6.** Mass spectrum of $[Fe_2(L2)_3](BF_4)_4$ 5
- **Figure S7.** Paramagnetic NMR spectrum of $[Ni_2(L2)_3](BF_4)_4$ 5
- **Figure S8.** Mass spectrum of $[Ni_2(L2)_3](BF_4)_4$ 6
- **Figure S9.** Mass spectrum of $[Ni_2(L2)_3](OAc)_4$ 6
- **Figure S10.** Photos of $[Fe_2(L2)_3](OAc)_4$ solutions over time 7
- **Figure S11.** UV-VIS spectra of $[Ni_2(L2)_3](OAc)_4$ over time 7
- **Figure S12.** Mass spectrum of $[Ni_2(L2)_3](OAc)_4$ after 12 hours in aqueous buffer 8
- **Figure S13.** MD snapshot of the helicate bound at the duplex terminus of the 3WJ 8
- **Figure S14.** $\pi$-stacking interactions of the L1 cylinder bound in the 3WJ 9
- **Figure S15.** MD snapshot of the mesocate bound inside a frayed 3WJ 9
- **Figure S16.** MD snapshots of the mesocate bound in the 3WJ 10
- **Figure S17.** Representative PAGE competition gels 10
- **Figure S18.** MD snapshot of the helicate bound inside a partially closed 4WJ 11
- **Figure S19.** Zoom-in MD snapshot of the mesocate inside the rhombus-shaped 4WJ cavity 11
- **Figure S20.** MD snapshots of dsDNA binding 11

Experimental Methods 12
- Synthesis of ligand L2 12
- Synthesis of $[Fe_2(L2)_3](BF_4)_4$ 12
- Synthesis of $[Ni_2(L2)_3](BF_4)_4$ 12
- Synthesis of $[Ni_2(L2)_3](OAc)_4$ 12
- DNA Sequences 13
- Polyacrylamide Gel Electrophoresis (PAGE) 13
- PAGE Competition Experiments 13
- UV-VIS Spectroscopy 14
- Circular Dichroism (CD) 14
- Linear Dichroism (LD) 14

Molecular Dynamics Simulations 14
- Parameterisation of the Compounds 14
- Parameterisation of DNA 14
- Simulations 14

References 15

### SUPPLEMENTARY DATA

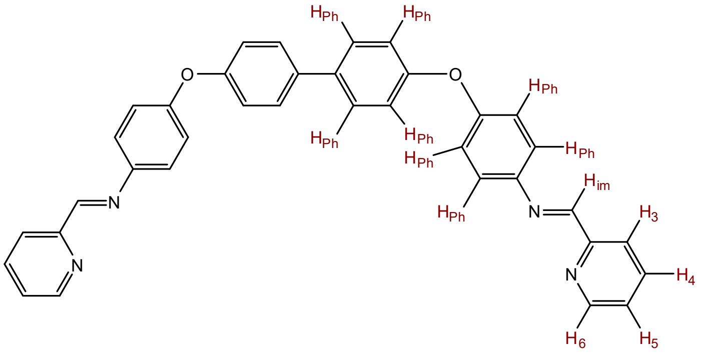


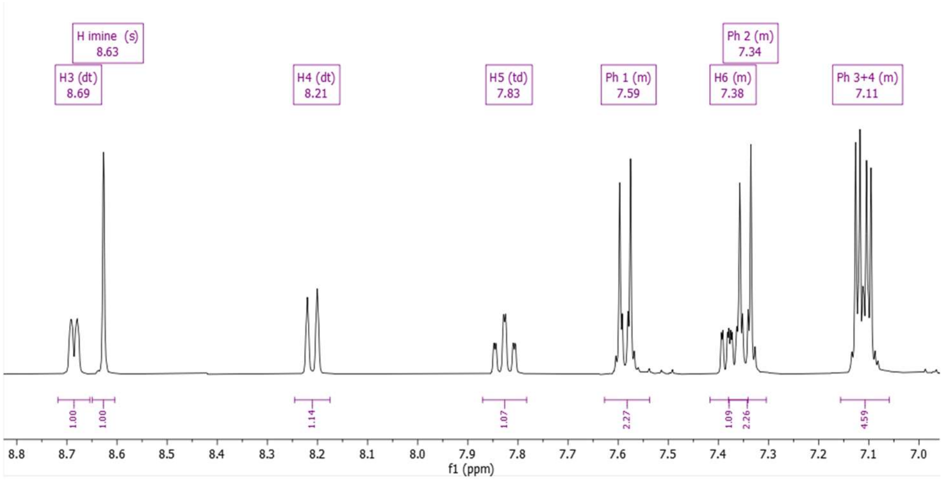


**Figure S1.** $^{1}H$ NMR spectrum (400 MHz, $CD_2Cl_2$) of ligand L2 showing assignments.

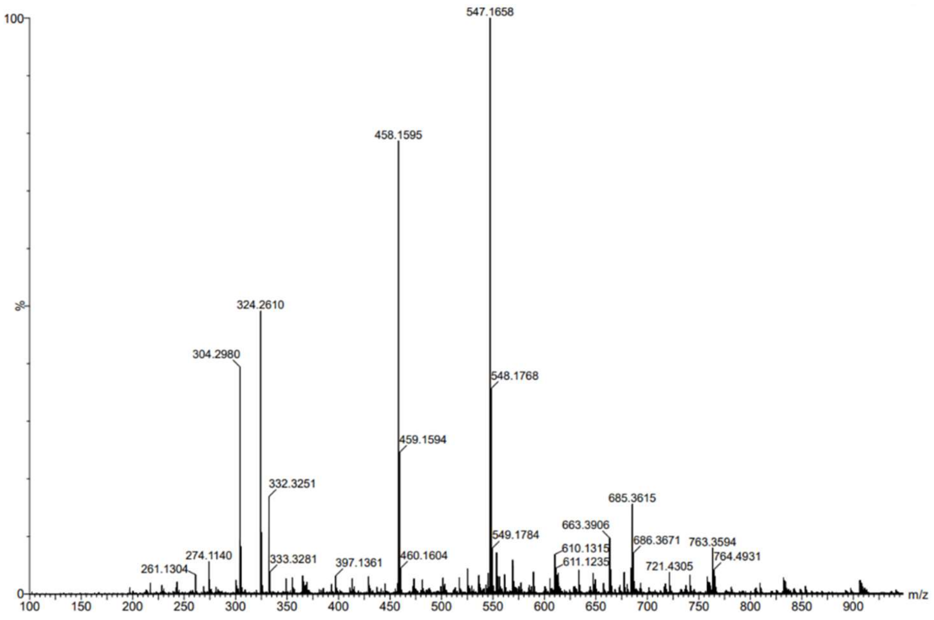


**Figure S2.** ESI Mass Spectrum of L2 (ESI+): m/z 547 ($[L2 + H]^+$, calc 547.21), m/z 458 ($[C_{30}H_{23}N_3O_2 + H]^+$ calc 458.19; corresponds to hydrolysis of one imine bond).

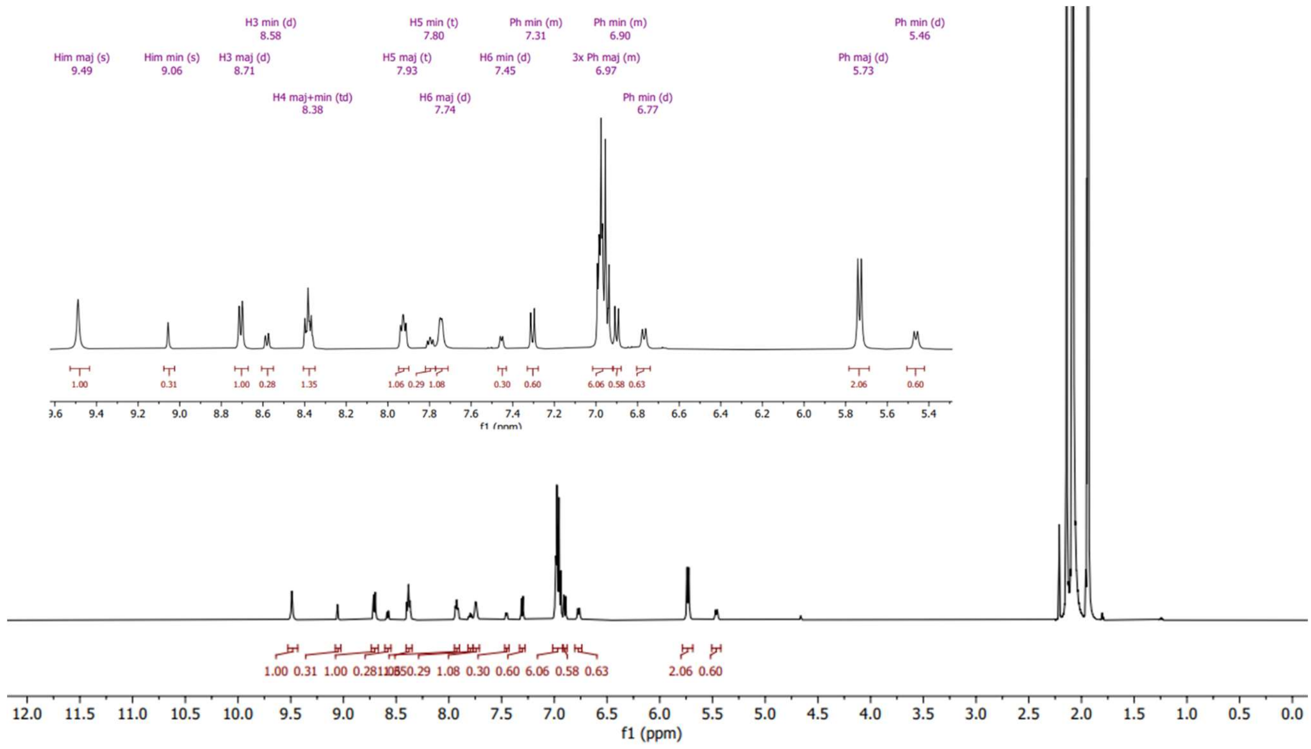


**Figure S3.** [1]H NMR spectrum (500 MHz, acetonitrile-d3) of $[Fe_2(L2)_3][BF_4]_4$ showing assignments for both the major (maj) and minor (min) species.

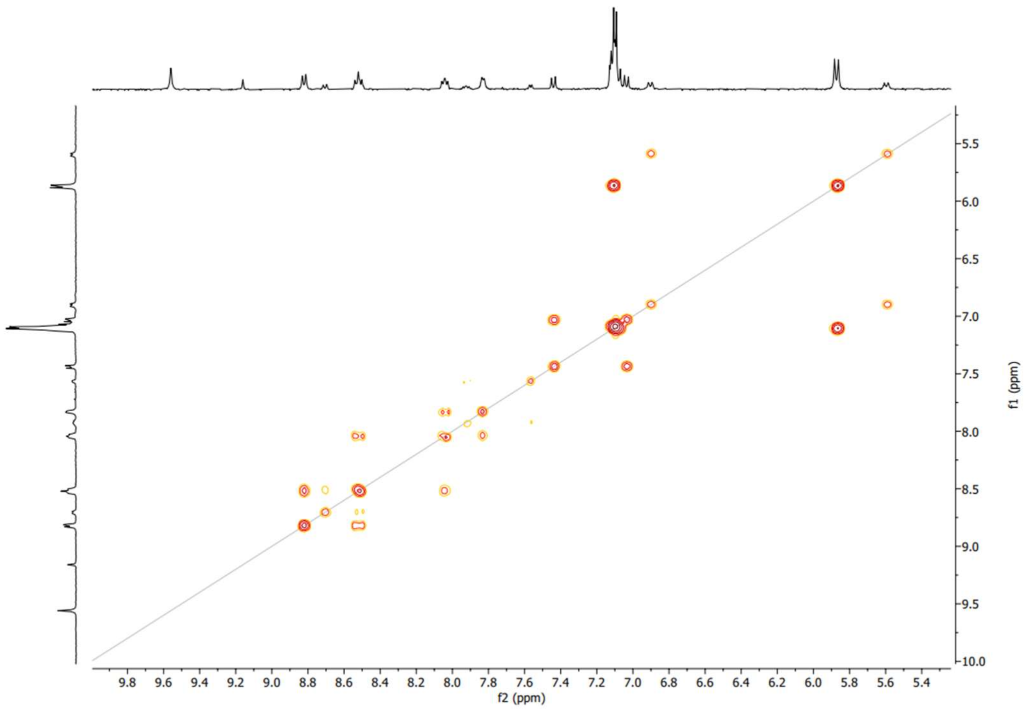


**Figure S4.** COSY NMR of $[Fe_2(L2)_3](BF_4)_4$ (400 MHz, acetonitrile-d3).

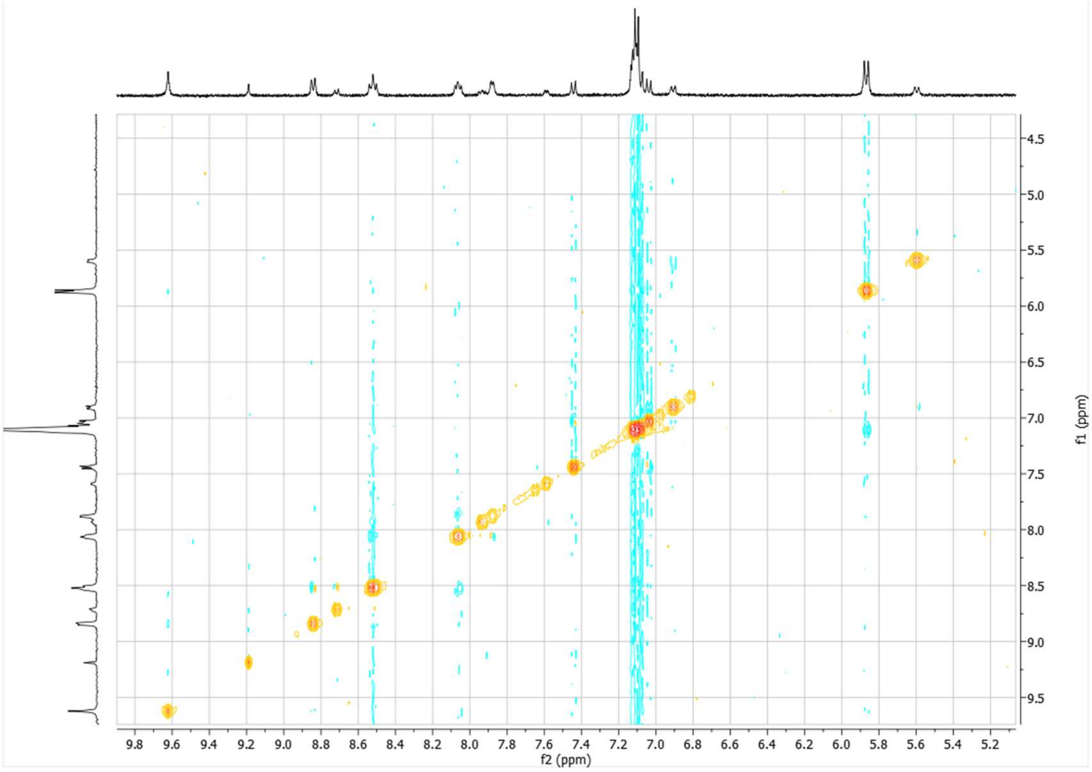


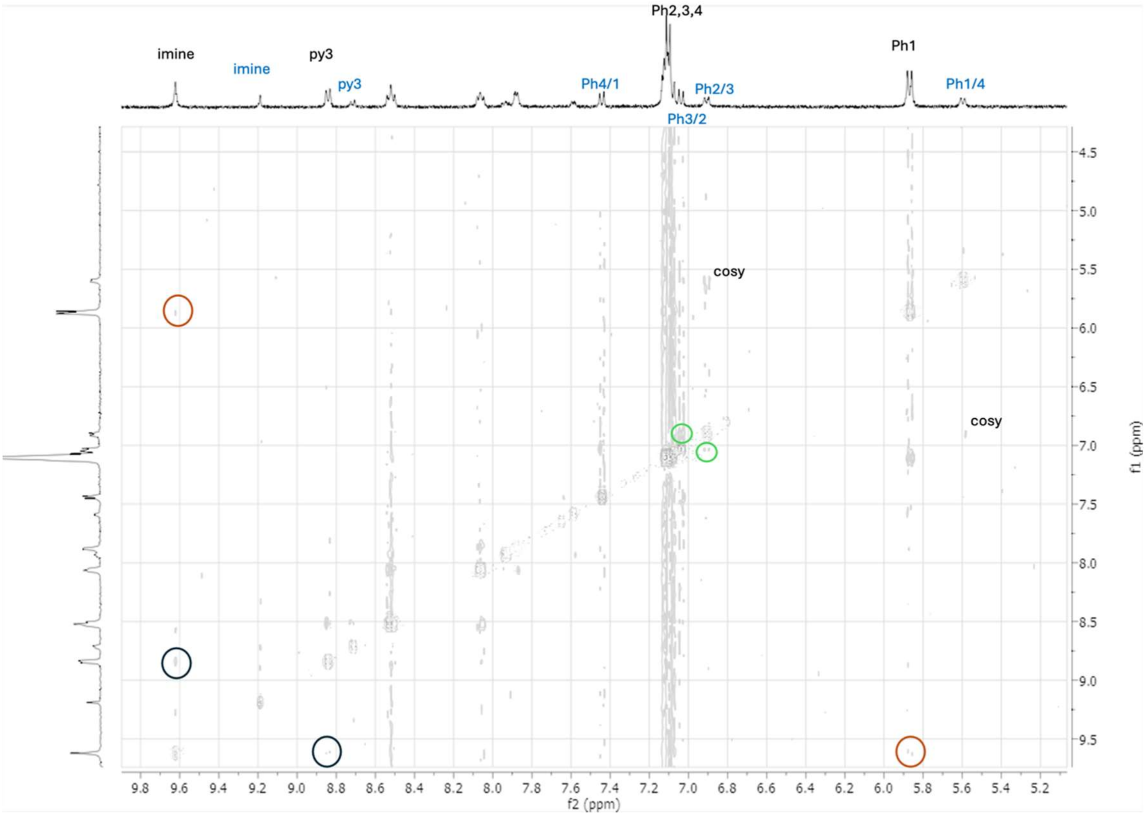


**Figure S5.** NOESY NMR of $[Fe_2(L2)_3](BF_4)_4$ (500 MHz, acetonitrile-d3). Three key NOE signals are circled.

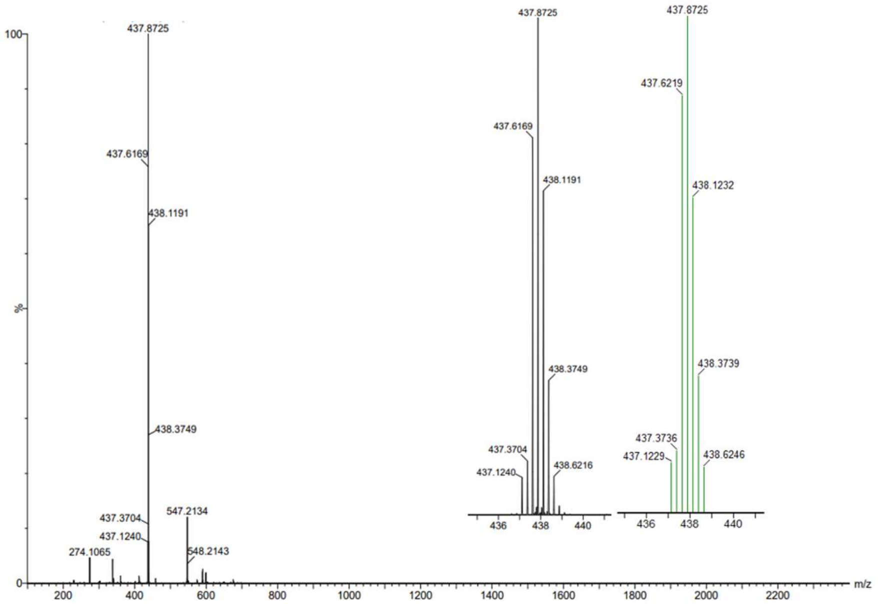


**Figure S6.** Electrospray mass spectrum of $[Fe_2(L2)_3](BF_4)_4$ ($ESI^+$): m/z 437 ($[Fe_2(L2)_3]^{4+}$, calc 437.87), m/z 547 ($[L2 + H]^+$, calc 547.21). Blow up shows the observed 437 peak (left) and the corresponding prediction (right).

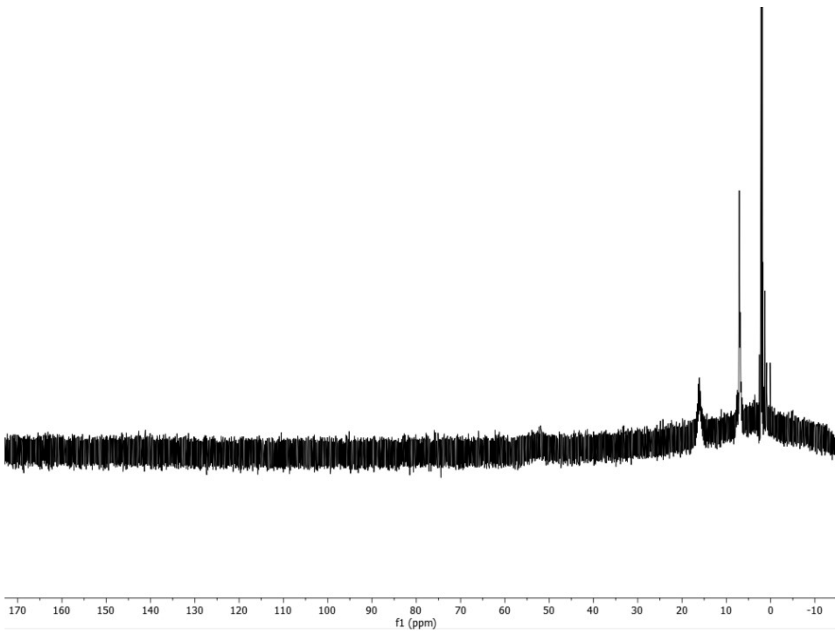


**Figure S7.** Paramagnetically shifted $^1H$ NMR $[Ni_2(L2)_3](BF_4)_4$ (300 MHz, $CD_3OD$)

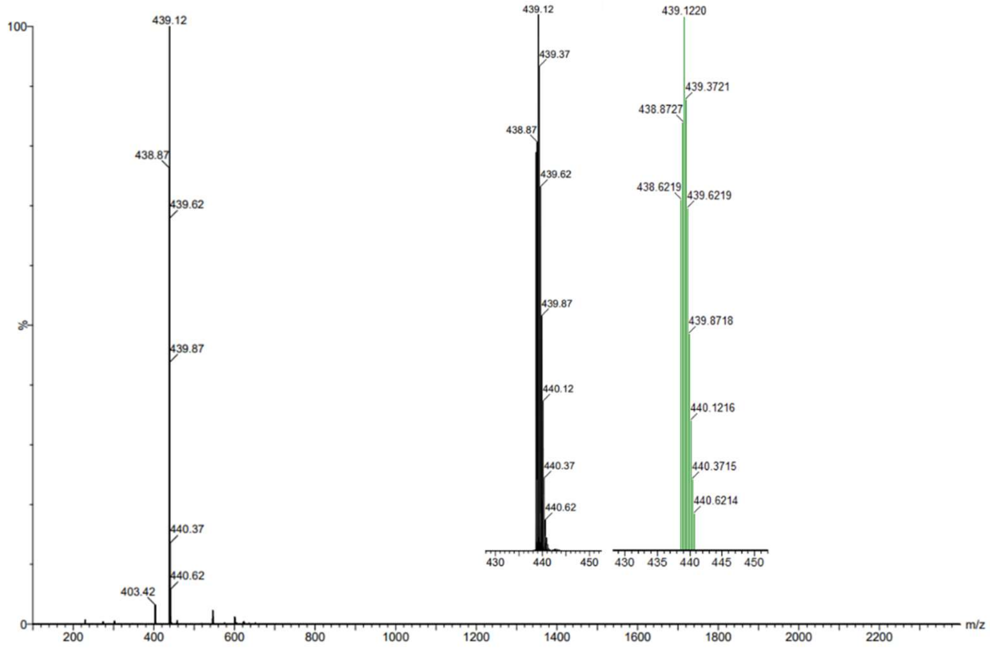


**Figure S8.** Electrospray mass spectrum of $[Ni_2(L2)_3](BF_4)_4$ ($ESI^+$): m/z 439 ($[Ni_2(L2)_3]^{4+}$, calc 439.12), 403 ($[L2+K]^+$, 5%) . Blow up shows the observed 439 peak (left) and the corresponding prediction (right).

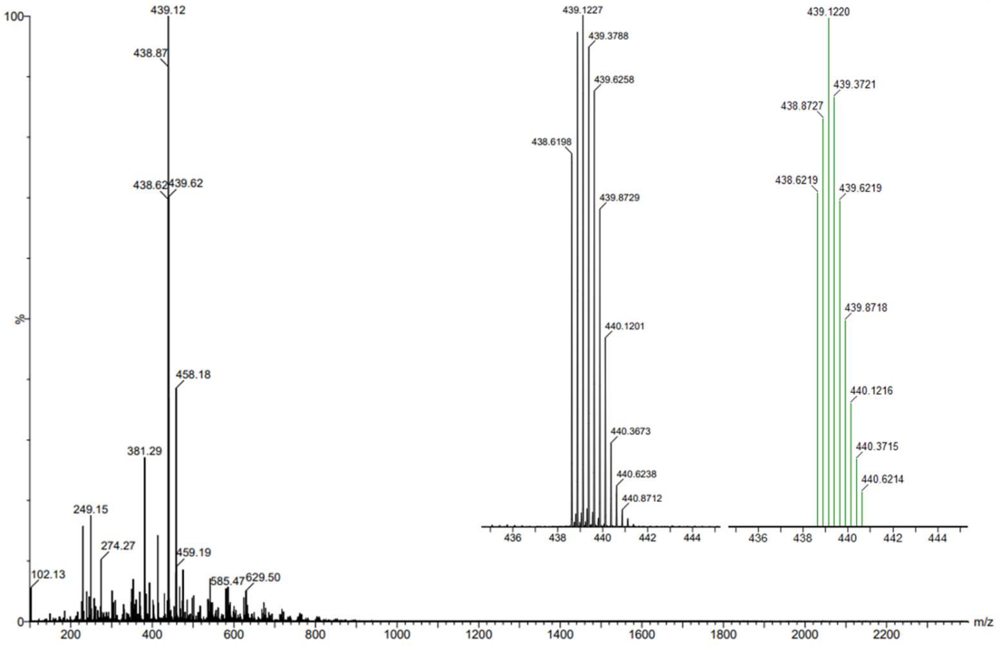


**Figure S9.** Electrospray mass spectrum of $[Ni_2(L2)_3](OAc_4)_4$. Blow up shows the observed 439 peak (left) and the corresponding prediction (right). ($ESI^+$): m/z 439 ($[Ni_2(L2)_3]^{4+}$, calc 439.12).

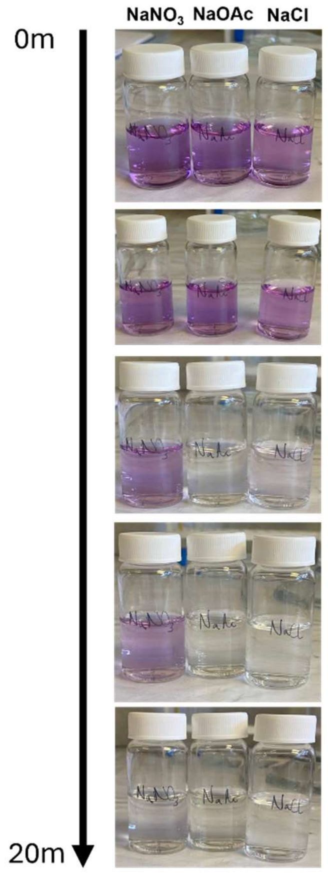


**Figure S10.** Photos of solutions of $[Fe_2(L2)_3](OAc)_4$ in $NaNO_3$ (left), NaOAc (middle) and NaCl (right). After 20 minutes, all solutions have lost their purple colour due to degradation of the cylinder.

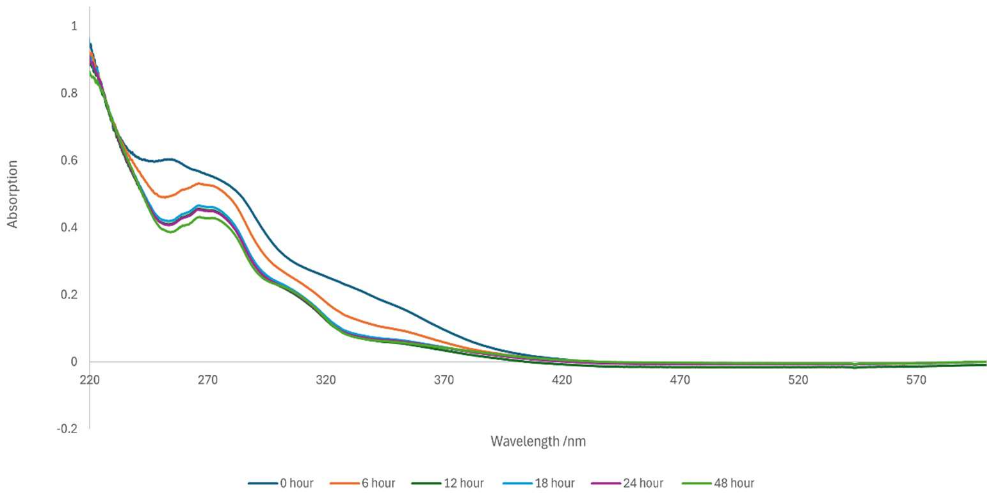


**Figure S11.** UV-VIS absorption spectra of $[Ni_2(L2)_3](OAc)_4$ over time (20 µM in water).

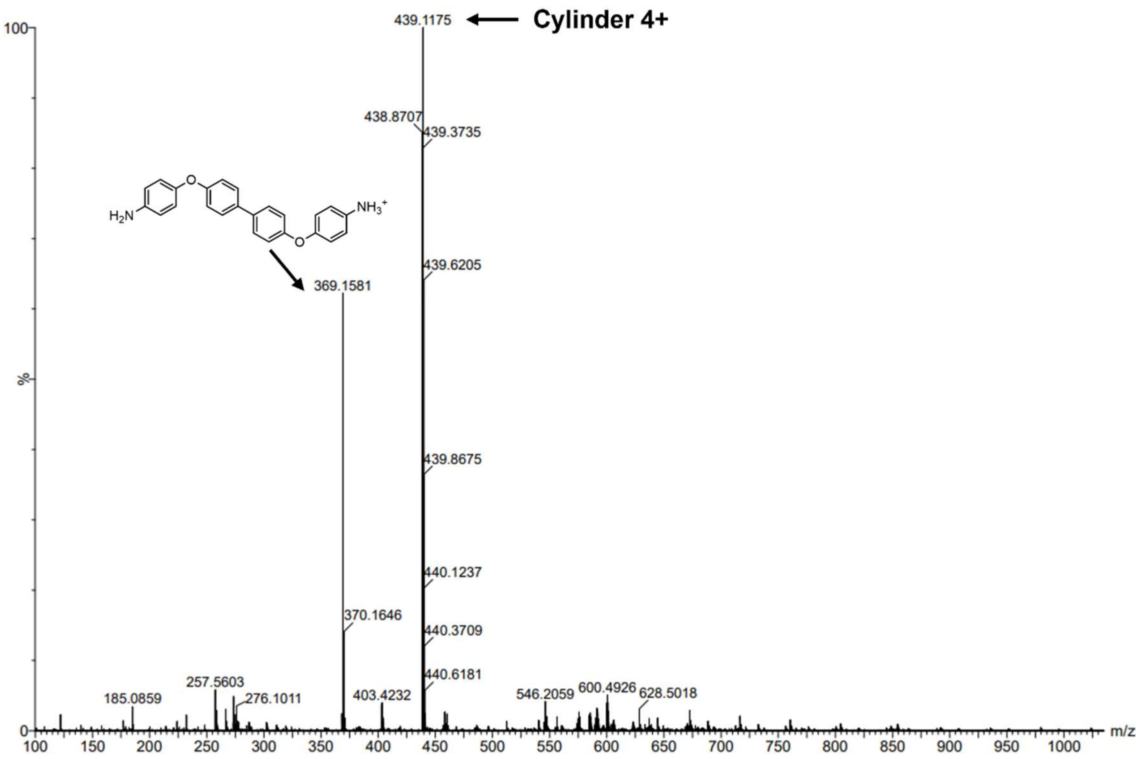


**Figure S12.** Mass Spectrum of $[Ni_2(L2)_3](OAc)_4$ cylinder in TBN buffer after 12 hours. m/z 439 ($[Ni_2(L2)_3]^{4+}$, calc 437.87),369 ($C_{24}H_{20}N_2O_2 + H^+$, calc 369.16).

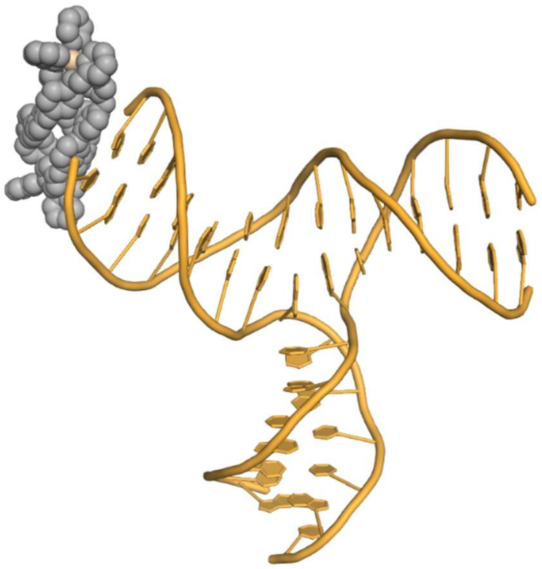

**Figure S13.** MD snapshot of the L2 helicate (ΔΔ enantiomer) bound at the duplex terminus of the 3WJ.

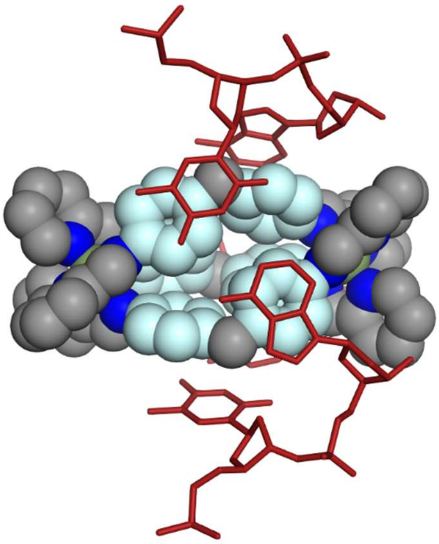

**Figure S14.** The crystal structure for the ΛΛ L1 cylinder with the 3WJ. Only the branchpoint base pairs are shown (red) and the phenyl moieties on the cylinder involved in π-stacking are highlighted in light blue. As seen for the L2 cylinder in Figure 4B, the aromatic rings across two adjacent ligands orient themselves coplanar to stack with the branchpoint base pairs.

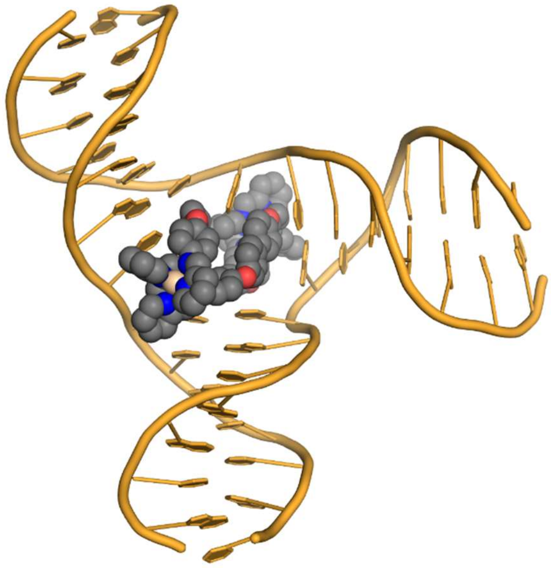

**Figure S15**. MD snapshot image of the ΛΛΔ mesocate isomer bound inside a frayed 3WJ cavity. Hydrogens have been omitted for clarity.

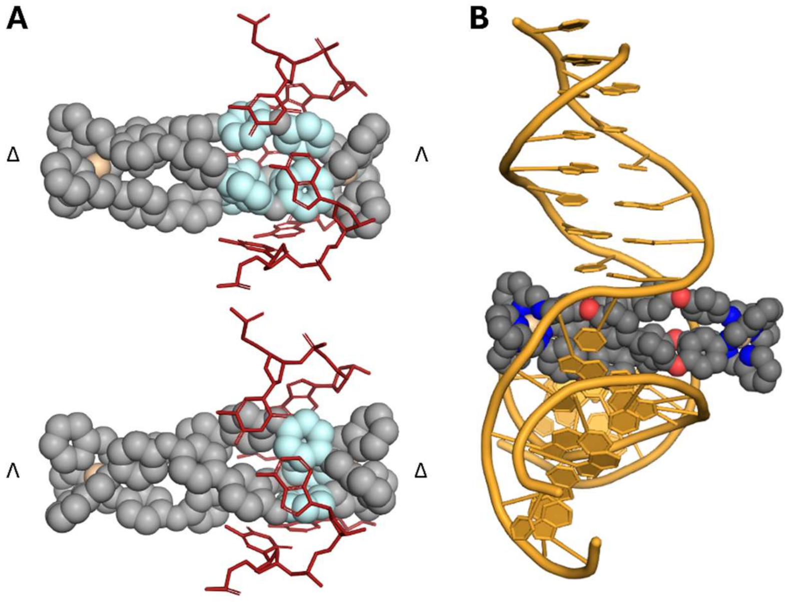


**Figure S16.** A) Close-up MD snapshot images of the ΛΔΔ mesocate isomer in its Λ-bound position (top) and Δ-bound position (bottom) in the 3WJ. The aromatic surfaces involved in face-face pi stacking are coloured in light blue. Only the branchpoint DNA bases are shown (red). B) MD snapshot of the same isomer bound inside the 3WJ. In this particular simulation, the mesocate was placed with the Λ end bound in the cavity but moved over the course of the simulation towards binding with Δ end, placing the mesocate much more centrally in the cavity as opposed to the binding modes seen in Fig. S16A. Hydrogens have been omitted in all images for clarity.

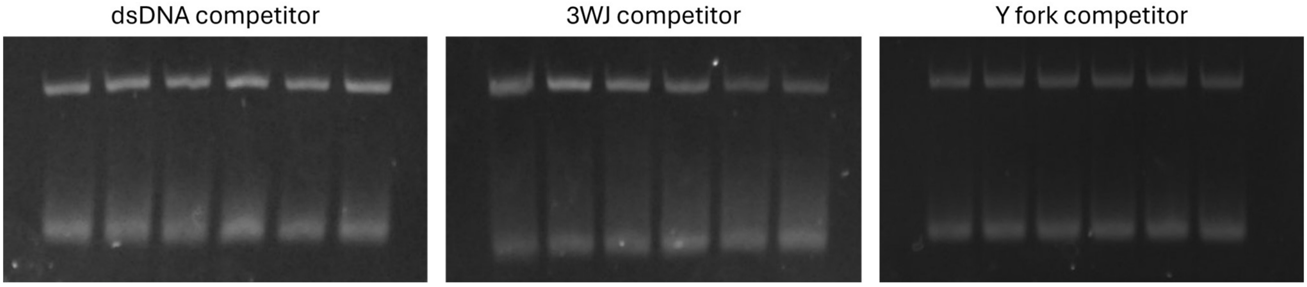


**Figure S17.** Representative PAGE gels in the competition assay. Each lane contains 2 μM 4WJ DNA and 6 μM Ni L2 cylinder. Increasing amounts (0, 0.5, 1, 2, 3, 4 equiv.) of competitor DNA (dsDNA, 3WJ or Y fork) are added from left to right.

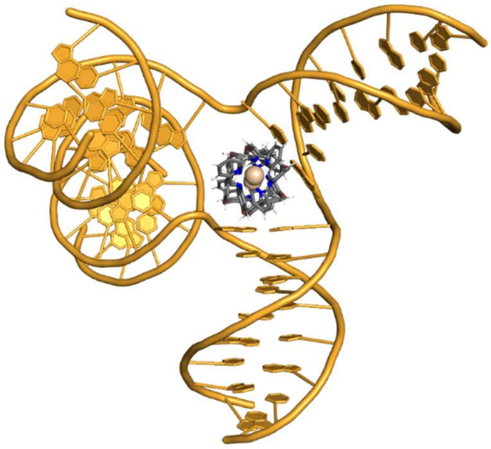

**Figure S18.** MD snapshot of a metastable conformation of the 4WJ with the Ni L2 ΛΛ helicate bound. The cylinder is bound in a partially closed 4WJ cavity, where two duplex arms are able to stack coaxially, leaving a 3WJ-like cavity space available to the cylinder, with base pair-cylinder interactions on 2 out of 3 sides.

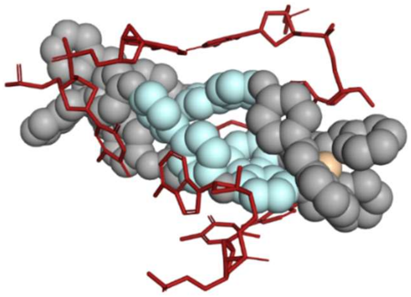

**Figure S19**. Close-up MD snapshot image of the ΛΔΔ mesocate isomer bound inside a rhombic-shaped 4WJ cavity. The aromatic surfaces involved in face-face pi stacking are coloured in light blue. Only the branchpoint DNA bases are shown (red). Hydrogens have been omitted for clarity.

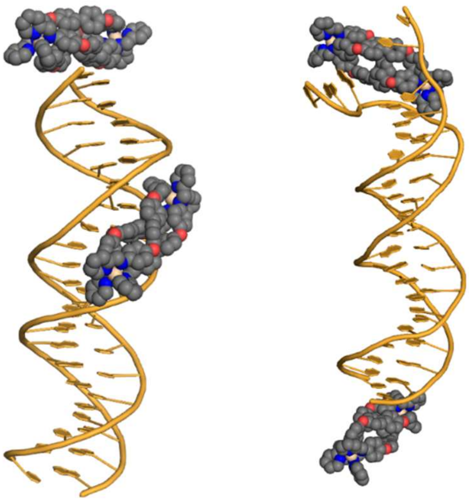

**Figure S20.** MD snapshots of 2 mesocates (left) and 2 helicates (right) binding to dsDNA. The mesocate snapshot shows minor groove binding and terminus binding. The helicate snapshot shows terminus binding at both ends, and where one end is frayed.

### EXPERIMENTAL METHODS

All solvents, NMR solvents, chemical reagents and buffer components, were purchased from Fischer Scientific, VWR chemicals or Sigma Aldrich and used without further purification. DNA oligomers were purchased dry and HPLC purified from Sigma Aldrich and used without further purification. Nickel cylinder $[Ni_2(L1)_3]Cl_4$ was prepared as described previously [1]. Electrospray ionisation (ESI) mass spectrometry characterisation was carried out on a Waters SYNAPT-G2-S in positive ion mode. 1H NMR studies were carried out on AVIII 300 (300 MHz), AVANCE NEO400 (400 MHz) and AVANCE NEO500 (500 MHz) Bruker spectrometers. Milli-Q (18.2 MΩ) water was used in all biophysical experiments.

**Synthesis of Ligand L2**

Ligand (L2) was synthesised as follows: 4,4'-(1,1'-biphenyl-4,4'-diyldioxy)dianiline (250 mg, 0.679 mmol) was dissolved in ethanol (25 mL) and pyridine-2-carboxaldehyde (0.44 mL, 4.67 mmol, excess) and a couple of drops of acetic acid were added dropwise. The solution was heated under reflux for 24 hours and the off-white precipitate that formed was collected by filtration, washed with cold ethanol and then dried *in vacuo.* Yield: 299 mg (80 %)

1H NMR (400 MHz, $CD_2Cl_2$) δ 8.69 (br d, J = 4.7 Hz, 1H, $H_{6py}$), 8.63 (s, 1H, $H_{imine}$), 8.21 (br d, J = 7.8 Hz, 1H, $H_{3py}$), 7.83 (td, J = 7.6, 1.7 Hz, 1H, $H_{4py}$), 7.59 (d, J = 8.6 Hz, 2H, $H_{Ph}$), 7.38 (ddd, J = 7.5, 4.7, 1.2 Hz, 1H, $H_{5py}$), 7.35 (d, J = 8.9 Hz, 2H, $H_{Ph}$), 7.12 (m comprising two overlapping d, J~ 8.8 Hz, 4H, 2x $H_{Ph}$)

MS (ESI+): *m/z* 547 ($[M+H]^+$), 458 ($[C_{30}H_{23}N_3O_2]^+$; hydrolysis of imine bond).

**Synthesis of $[Fe_2(L2)_3](BF_4)_4$**

Ligand L2 (30 mg, 0.055 mmol) was dissolved in acetonitrile (5 mL) and heated to 50ºC. Iron(II) tetrafluoroborate hexahydrate (12.4 mg, 0.037 mmol) was dissolved in acetonitrile (5 mL) and added to the ligand mixture dropwise with stirring. The reaction mixture was then stirred and heated under reflux (80ºC) for 4 hours. The purple precipitate was collected by vacuum filtration and redissolved in the minimum volume of acetonitrile. Finally, the purple product was precipitated by addition of cold diethyl ether (50 mL) and collected by vacuum filtration Yield: 26 mg (68 %)

1H NMR (500 MHz, CD3CN) δ 9.49 (s, 7H, $H_{imine}$ maj), 9.06 (s, 2H, $H_{imine}$ min), 8.71 (d, J = 7.6 Hz, 7H, $H_{3py}$ maj), 8.58 (d, J = 7.6 Hz, 2H, $H_{3py}$ min), 8.38 (m, comprising two overlapping t, J ~ 7.6Hz, 9H, $H_{4py}$ maj + $H_{4py}$ min), 7.93 (br t, J = 6.6 Hz, 7H, $H_{5py}$ maj), 7.80 (br t, J = 6.6 Hz, 2H, $H_{5py}$ min), 7.74 (br d, J = 5.3 Hz, 7H, $H_{6py}$ maj), 7.45 (br d, J = 5.5 Hz, 2H, $H_{6py}$ min), 7.31 (d, J = 8.6 Hz, 4H, $H_{Ph}$ min), 6.98 (m, 42H, 3x $H_{Ph}$ maj), 6.90 (d, J = 7.6 Hz, 4H, $H_{Ph}$ min), 6.77 (br d, J = 8.2 Hz, 4H, $H_{Ph}$ min), 5.73 (d, J = 8.3 Hz, 14H, $H_{Ph}$ maj), 5.46 (br d, J = 8.3 Hz, 4H, $H_{Ph}$ min).

MS (ESI+): m/z 437 ($[Fe_2(L2)_3]^{4+}$, 100%), m/z 547 ($C_{36}H_{26}O_2N_4^+$, 10%), m/z 274 ($C_{18}H_{13}ON_2^+$, 5%).

**Synthesis of $[Ni_2(L2)_3](BF_4)_4$**

Ligand L2 (30 mg, 0.055 mmol) was dissolved in acetonitrile (5 mL) and heated to 50ºC. Nickel(II) chloride tetrahydrate (8.8 mg, 0.037 mmol) was dissolved in acetonitrile (5 mL) and added to the ligand mixture dropwise with stirring. The reaction mixture was then heated (55ºC) with stirring for 2 hours. The precipitate was collected by vacuum filtration and redissolved in the minimum volume of acetonitrile. The yellow-orange product was precipitated by treating with 50 mL cold diethyl ether and collected by vacuum filtration. Yield: 28 mg (93 %)

MS (ESI+): m/z 439 ($[Ni_2(L2)_3]^{4+}$, 100%).

**Synthesis of $[Ni_2(L2)_3][OAc]_4$**

The ligand (L2) (30 mg, 0.055 mmol) was dissolved in methanol (5 mL) which was heated to 50ºC. Nickel(II) acetate tetrahydrate (9.1 mg, 0.037 mmol) was dissolved in methanol (5 mL) and added to the ligand mixture dropwise with stirring. The reaction mixture was then heated at 55° C for 4 hours. The precipitate was collected by vacuum filtration and redissolved in the minimum volume of acetonitrile. Finally, the product was crashed out in cold diethyl ether and isolated by vacuum filtration Yield: 15.1 mg (38 %)

MS (ESI+): m/z 439 ($[M_2(L2)_3]^{4+}$, 100%), 458 ($[C_{30}H_{23}N_3O_2]^+$, 37%), 381 ($[C_{25}H_{20}N_2O_2]^+$, 27%), 274 ($[C_{18}H_{13}N_2O]^+$, 5%),

**DNA Sequences (5′ to 3′)**

DS-21 S1: CCTTCACGCGAACGTAATCCT
DS-21 S2: AGGATTACGTTCGCGTGAAGG

4WJ S1: GCTAGCTGATACGCTACG
4WJ S2: CGTAGCGTACGTTGGTGC
4WJ S3: GCACCAACGCGTCACTCC
4WJ S4: GGAGTGACGTCAGCTAGC

3WJ S1: CGGAACGGCACTCG
3WJ S2: CGAGTGCAGCGTGG
3WJ S3: CCACGCTCGTTCCG

Y fork S1: CGCACGTACGGAACGGCACTCGCTTGCTCG
Y fork S2: CGAGCAAGCGAGTGCAGCGTGGATACATGC

**Polyacrylamide Gel Electrophoresis (PAGE)**
20cm x 20cm large 12% PAGE gels were prepared by mixing 20 mL of 37.5:1 acrylamide/bis-acrylamide with 5 mL of 10X Tris-Boric acid buffer (890 mM, pH 8.3) and 25 mL of Milli-Q water. To this 400 µL of a 10% w/v ammonium persulfate solution in water and 40 µL of TEMED were added to initialise polymerisation. This was then immediately poured between 2 glass plates and a 20-well comb inserted at the top; this was then allowed to set for 1 hr before proceeding.

8.3cm x 7.3 cm mini 12% PAGE gels were prepared by mixing 6 mL of 37.5:1 acrylamide/bis-acrylamide with 1.5 mL of 10X Tris-Boric acid (TB) buffer (890 mM, pH 8.3) and 7.5 mL of Milli-Q water. To this 150 µL of a 10% w/v ammonium persulfate solution in water and 15 µL of TEMED were added to initialise polymerisation. This was then immediately poured between 2 glass plates and a 10-well comb inserted at the top; this was then allowed to set for 1 hr before proceeding.

The gel was then attached to the gel jacket and submerged in 1X TB buffer at the top and bottom. The wells were thoroughly flushed before loading of any sample. Samples were made up to 30 µL containing 1 µM of each DNA strand, 1X buffer (89 mM Tris, 89 mM Boric acid, 10 mM NaCl, pH 8.3), and the indicated ratio of complex. DNA, water, and buffer were mixed in solution before addition of the stated ratios of complex. Samples were then centrifuged and incubated at 37° C for 1 hr. 7.5 µL of 50% v/v glycerol was then added to each sample (10% v/v final concentration) and the sample was then centrifuged, mixed, and 10 µL pipetted into the wells on the gel. The gels were run at 140 V for 35 mins (mini gels) or 60 mins (big gels) in 1X TB running buffer. The gel was then removed from the plates and stained using SYBR™ Gold Nucleic Acid Gel Stain (Thermofisher scientific) in 1X TB buffer for at least 30 minutes before imaging on an AlphaImager™ UV transilluminator (Alpha Innotech) with 302 nm excitation.

**PAGE Competition Experiments**
8.3cm x 7.3 cm mini 12% PAGE gels were prepared as described previously in this document. The gel was then attached to the gel jacket and submerged in 1X TB buffer at the top and bottom. The wells were thoroughly flushed before loading of any sample. Samples were made up to 20 µL containing 2 µM of fluorescently labelled 3WJ (containing a 5′ FAM label on strand S1), 1X TBN buffer (89 mM Tris, 89 mM Boric acid, 50 mM NaCl, pH 8.3), 6 µM $[Ni_2(L2)_3](OAc)_4$ (3 equiv.), and the indicated ratio of competitor DNA. DNA, water, and buffer were mixed in solution before addition of the stated ratios of complex. Samples were then centrifuged and

incubated at room temperature for 15 mins. 5 µL of 50% v/v glycerol was then added to each sample (10% v/v final concentration) and the sample subsequently centrifuged, mixed, and 10 µL pipetted into the wells on the gel. The gels were run at 140 V for 35 mins in 1X TB running buffer. The gel was then removed from the plates and rinsed in deionised water for 5 minutes before imaging on an AlphaImager™ UV transilluminator (Alpha Innotech) with 302 nm excitation. ImageJ was used to quantify the intensity of the gel bands [2]. The intensities of the 3WJ bands were measured as a fraction of the total lane intensity (ssDNA band intensity + 3WJ band intensity) and normalised to the lane containing no competitor DNA. All competitor ratios were measured as the average of 3 independent samples and plotted with standard deviation error bars.

**UV-VIS Spectroscopy**

Absorbance was recorded between 200-800 nm (1 nm bandwidth, 600 nm/min) in a Cary5000 UV-Vis-NIR Spectrophotometer (Agilent Technologies, Inc.) equipped with a multi-cell holder, using a 1 cm path length, masked quartz cuvette. In all cases, each spectrum was zeroed and a baseline correction recorded for each condition.

**Circular Dichroism (CD)**

CD spectra were recorded on a Chirascan+ spectrophotometer (Applied Photophysics Ltd.). The samples were scanned in a 1cm path length, quartz, masked cuvette from 600 nm to 200 nm (1 nm SBW, 0.5 s averaging time). Each condition was repeated 3 times (fresh sample each time). Samples were prepared in 10 mM Na cacodylate and 100 mM NaCl, with ctDNA concentration of 200 µM in DNA base pairs.

**Linear Dichroism (LD)**

LD spectra spectra were recorded on a Chirascan+ spectrophotometer (Applied Photophysics Ltd.) using the LD accessory. Samples were scanned in a quartz couette cell with an angular gap of 0.25 mm, giving an overall path length of 0.5 mm. The couette cell was rotated at 40 revolutions per second with a 3 minute incubation time at a rotation of 3 revolutions per second at each addition. A baseline correction was recorded by scanning the DNA alone with no rotation. Titrations began at a volume of 150 µL and did not exceed 250 µL. Each condition was repeated 3 times (fresh sample each time). Samples were prepared in 10 mM Na cacodylate and 100 mM NaCl, with ctDNA concentration of 200 µM in DNA base pairs.

### MOLECULAR DYNAMICS SIMULATIONS

**Parameterisation of the Compounds**

Parameters for the coordination bonds were calculated using the MCPB.py pipeline with Gaussian16 at the ωB97XD/DEF2-SVP level of theory to include dispersion [3, 4]. The output coordinate and topology files were then converted to GROMACS format (.top and .gro) using ParmEd (https://github.com/ParmEd/ParmEd). Coordinates for the mirror image enantiomers were generated using the invert chirality function in Avogadro [5].

**Parameterisation of DNA**

The PDB file for the 25mer B-DNA consisting of 2 strands ($A_{25}$ and $T_{25}$) was generated using NAB (nucleic acid builder) in AmberTools [6]. The 3WJ structure was adapted from PDB 1F44 [7], as described previously [8]. The 4WJ structure was taken from the 1XNS PDB crystal structure [9], and the strand lengths shortened as described previously [8]. All DNA was parameterised using the AMBER forcefield parmbsc1 [10].

**Simulations**

In simulations with the 1XNS 4WJ, the compounds were placed either inside or outside but close to the open cavity of the structure. In all simulations of 3WJ, the compounds were similarly placed either inside or outside but close to the cavity. In all simulations with the B-DNA, multiple compounds were placed within 1 nm distance of the DNA. In all simulations, DNA was placed with the compounds in a dodecahedral box with periodic boundary conditions. MD preparation steps were carried out using GROMACS software as described previously [8, 11-14]: All systems were solvated in water using the TIP3P model and neutralised with $Na^+$ ions. Additional $Na^+$ and $Cl^-$ ions were added to reach a NaCl concentration of 50 mM. Initial minimisation was carried to at least 500 kJ/mol/nm or 50000 steps followed by heating and NVT equilibration for 1000 ps using

V-rescale modified Berendsen thermostat, coupling the cylinder with the DNA at 310 K. All simulations use 2 fs time step and Parrinello-Rahman pressure coupling and PME electrostatics at 1.0 nm cutoff. All simulations were run on the BlueBEAR cluster at U. Birmingham using GROMACS software. After the simulations had finished, the trajectories were processed in GROMACS to remove periodic boundary conditions, translations and rotations, and visualised in PyMOL [15].
